# CK1δ homeostasis by activity-dependent shuttling and degradation of orphan kinase

**DOI:** 10.1101/2023.02.13.528286

**Authors:** Fidel E. Serrano, Daniela Marzoll, Bianca Ruppert, Axel C. R. Diernfellner, Michael Brunner

## Abstract

Casein kinase 1δ (CK1δ) is a simple monomeric enzyme involved in the regulation of a variety of functions, including signal transduction, the circadian clock, and the cell cycle. Although CK1δ is targeted by the ubiquitin ligase APC/C^Cdh1^ is not understood how CK1δ expression is regulated to support its multiple functions. Here, we show that kinase activity controls CK1δ homeostasis by coordinating two competing processes: export from the nucleus to ensure distribution of CK1δ between its assembly partners, and proteasomal degradation of unassembled CK1δ in the nucleus to keep the amount of active, potentially deleterious orphan kinase low. During mitosis, CK1δ is released from centrosomes and stabilized by (auto)phosphorylation to preserve it for the subsequent G1 phase.

**Teaser:** Competitive nuclear export and nuclear degradation of active CK1δ ensure efficient partner interaction and keep unassembled kinase levels low.

## Introduction

Casein kinase 1 (CK1) is conserved across the eukaryotic lineage (*1*) with homologs in yeast (*2*), fungi (*3*), plants (*4*), and animals (*5*). CK1 family members share a highly conserved structured catalytic domain and an unstructured non-conserved C-terminal tail, which can be (auto)phoshorylated and inhibit the kinase (*6, 7*). Several autophosphorylation sites in the C-terminal tail of CK1 have been identified (*8*). CK1 members serve key roles as effectors of various cellular signalling pathways (*9, 10*)

Of the seven vertebrate CK1 isoforms (α, β, γ1, γ2, γ3, δ and ɛ) CK1δ, encoded by *CSNK1D* in humans, has a function in the circadian clock (*5, 11, 12*), cytoskeletal organization (*13, 14*), cell cycle (*15, 16*), and development (*17–19*). CK1δ shuttles between the cytosol and the nucleus and kinase activity affects its subcellular localization (*20*). In the context of developmental regulation, CK1δ has been shown to phosphorylate clients in the Wingless (*17*), Hedgehog (*18*), and Hippo pathways (*19*). CK1δ deregulation has been implicated in various diseases including neurological and metabolic pathologies and cancer (*9, 10*).

A fraction of CK1δ is recruited by Period (PER) proteins of the circadian clock, which constitute together with Cryptochromes (CRY) crucial elements of the circadian pacemaker (*5, 21–23*). In the course of a day, PERs are progressively phosphorylated by bound CK1δ and the CRY-PER-CK1δ complex forms a functionally conserved phosphorylation-based module to measure time on a circadian scale (*12, 24, 25*). Phosphorylation of two competing sites in PER, the β-TrCP and FASP sites, are major determinants of PER stability and circadian period length (*11, 26*). Along with regulating PERs’ stability and nuclear localization (*27*), CK1δ contributes to the negative feedback of PERs and CRYs on their transcription factor BMAL1-CLOCK by phosphorylating CLOCK and thereby displacing BMAL1-CLOCK from the DNA (*28–30*).

CK1δ contains a centrosomal localization signal that extends from the C-terminal portion of the kinase domain into the disordered C-terminal tail and a substantial fraction of CK1δ is associated with the centrosomes (*31*). CK1δ is involved in the cell cycle progression via Wee1 phosphorylation and inhibition resulting in enhanced entry into mitosis (*16*). During mitosis, CK1δ has been shown to phosphorylate tubulin regulating microtubule polymerization and stability of the spindle pole apparatus (*13, 32*). CK1δ is targeted for proteolysis via the cell cycle-specific ubiquitin ligase anaphase-promoting complex/cyclosome with its coactivator FZR1/CDH1 (hereafter APC/C^Cdh1^) (*33*).

Despite CK1δ being involved in a variety of cellular processes, there are open questions as to how CK1δ is regulated. CK1δ and its family members are monomeric enzymes that lack regulatory subunits and, unlike many other kinases, are not regulated by phosphorylation of their activation loop. Autophosphorylation of the C-terminal tail inhibits kinase activity (*6, 8*), but in the cell the kinase is maintained in an active dephosphorylated state, such that a physiological role for autoinhibition remains unknown.

Here we show that CK1δ bound to its interaction partners (e.g. centrosomes and PER2) is stable, whereas orphan CK1δ is rapidly turned over by proteasomal degradation in the nucleus. Degradation is facilitated by kinase activity, which is furthermore required for nuclear export of CK1δ. Autophosphorylation of the CK1δ tail or partially redundant phosphorylation by unknown kinase(s) stabilizes orphan kinase and leads to its accumulation in the nucleus. We also show that (auto)phosphorylated CK1δ accumulates in a cell cycle-dependent manner during G2/M. CK1δ is thereby inactivated and stabilized in late mitosis when APC/C^Cdh1^ becomes active and the kinase is no longer bound to centrosomes. Our data strongly suggest a physiological role for CK1δ (auto)phosphorylation in inhibiting and stabilizing orphan kinase during mitosis to maintain a CK1δ pool that can rapidly reassociate with centrosomes and other binding partners after cell division.

## Results

### Overexpressed active wild-type CK1δ is unstable

To study the regulation of CK1δ, we generated stable U2OS T-REx (U2OStx) cell lines expressing C-terminally FLAG-tagged wild-type CK1δ (U2OStx_CK1δ) and the kinase-dead version CK1δ-K38R (*34*) (U2OStx_CK1δ-K38R), respectively, when induced by doxycycline (DOX). Quantification of CK1δ RNA revealed that the induced transgenes were expressed at similar levels approximately 50-fold higher than the endogenous kinase (Fig. 1a). Immunoblot analyses revealed that FLAG-labeled CK1δ and CK1δ-K38R were already expressed at levels similar to endogenous CK1δ in the absence of DOX but were strongly overexpressed after DOX induction (Fig. 1b). However, CK1δ was expressed at a much lower level than the kinase-dead CK1δ-K38R, suggesting posttranscriptional regulation of the kinase in response to enzyme activity.

**Fig. 1.**
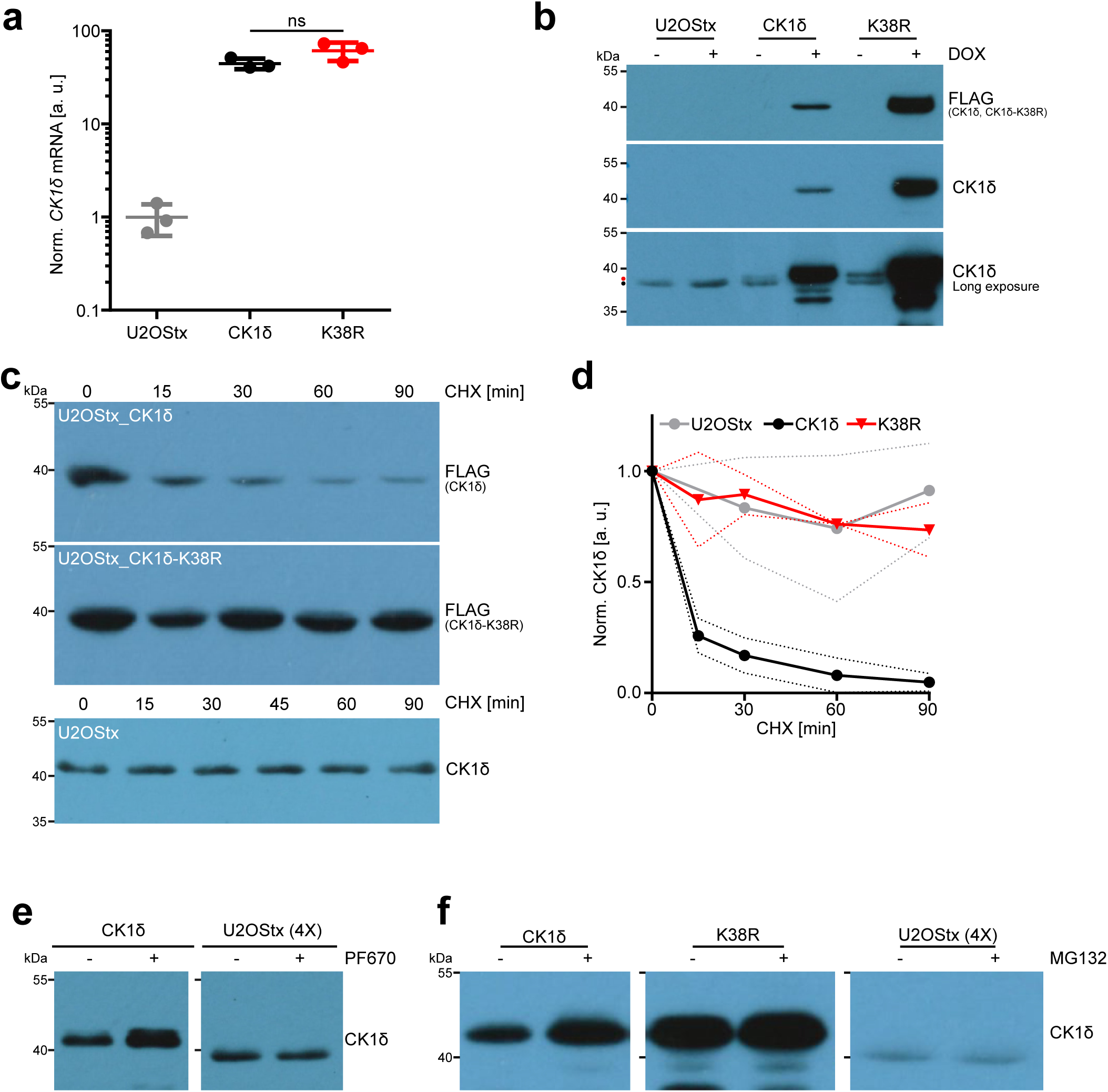
Overexpressed wild-type CK1δ is unstable. **a** mRNA expression levels of endogenous and transgenic CK1δ. RNA was prepared from U2OStx control cells and DOX-inducted stable U2OStx cell lines expressing CK1δ or CK1δ-K38R. Primers were specific to CK1δ recognizing both endogenous and transgenic CK1δ. CK1δ mRNA was quantified by RT-qPCR relative to GAPDH. **b** Protein expression levels of endogenous and transgenic CK1δ. U2OStx and stable U2OStx cell lines expressing inducible CK1δ and CK1δ-K38R were DOX-induced for 24 h. CK1δ protein levels were analyzed by immunoblotting with antibodies against either the FLAG epitope specific to the transgenic CK1δ or against CK1δ which recognizes both the endogenous and transgenic CK1δ. Black and red circles indicate endogenous and FLAG-tagged transgenic CK1δ, respectively **c** Overexpressed CK1δ is rapidly degraded, while CK1δ-K38R and endogenous CK1δ are stable over 90 min. U2OStx cell lines expressing CK1δ or CK1δ-K38R and U2OStx control cells expressing endogenous CK1δ were treated with CHX. Samples were collected at the indicated time points (CHX chase) and analyzed by immunoblotting against the FLAG epitope for transgenic CK1δ or against CK1δ for endogenous CK1δ. **d** Quantification of **c**. Results shown are averages of three independent experiments and are represented as the mean ± SD. The curve for the overexpressed CK1δ was used to calculate a half-life of about 6 min. **e** Kinase inhibition by PF670 results in accumulation of overexpressed CK1δ while having no effect on endogenous CK1δ. U2OStx and stable U2OStx cell lines overexpressing CK1δ were treated for 1 h with 1 µM PF670 or H_2_O vehicle control. 50 µg of U2OStx_CK1δ lysate (left panel) and 200 µg of U2OStx lysate (right panel) were loaded. CK1δ levels were analyzed by immunoblotting against CK1δ. **f** Proteasomal inhibition by MG132 stabilizes excess CK1δ and CK1δ-K38R while having no effect on endogenous CK1δ. Cells were treated for 4 h with 20 µM MG132 or DMSO (vehicle control). 50 µg of U2OStx_CK1δ (left panel) and U2OStx_CK1δ-K38R lysates (middle panel) and 200 µg of U2OStx lysate (right panel) were loaded. CK1δ levels were analyzed by immunoblotting against CK1δ.

We then tested whether the differences in kinase expression were due to protein turnover. Following a cycloheximide (CHX) chase, overexpressed CK1δ was rapidly degraded with a half-time of about 6 min, whereas CK1δ-K38R remained rather stable. Endogenous CK1δ, expressed in U2OStx control cells was also stable over the 90 min time course of the CHX treatment (Fig. 1c, d), in agreement with previous reports (*21, 30*).

Our observations suggest that overexpressed CK1δ is rapidly degraded depending on its enzymatic activity. We then tested how pharmacological inhibition of CK1δ by PF670462 (hereafter abbreviated as PF670), a well-characterized inhibitor of CK1δ and CK1ε (*35*), affects expression of endogenous and overexpressed CK1δ. PF670 treatment led to stabilization of overexpressed CK1δ whereas endogenous CK1δ abundance in U2OStx control cells was not affected (Fig. 1e), supporting that enzymatically active kinase is unstable when overexpressed.

CK1δ is a target of the APC/C^Cdh1^ ubiquitin ligase (*33*). To further characterize the degradation pathway of CK1δ, cells were treated with the proteasome inhibitor MG132. When the kinase was overexpressed, MG132 treatment led to a substantial increase in CK1δ abundance indicating that a large fraction of CK1δ synthesized from the overexpressed RNA was rapidly degraded in the absence of MG132 (Fig. 1f, left panel). MG132 treatment also stabilized CK1δ-K38R but to a lesser extent (Fig. 1f, middle panel), suggesting that kinase activity accelerates degradation of overexpressed CK1δ. When U2OStx control cells were treated with MG132, endogenous CK1δ levels did not increase (Fig. 1f, right panel). These data suggest that translation of the relatively low amounts of endogenous RNA was insufficient to significantly increase kinase levels within the 1 h and 4 h treatment windows with PF670 and MG132, respectively.

### Phosphorylation stabilizes CK1δ during G2/M

CK1δ contains (auto)phosphorylation sites in its C-terminal tail and phosphorylation at these sites leads to inhibition of the kinase(*8*). However, phosphorylated CK1δ does not accumulate in detectable amounts in the cell. Rather, CK1δ undergoes futile cycles of (auto)phosphorylation and dephosphorylation (*6*). Therefore, the physiological significance of phosphorylation-induced inhibition is unclear. We asked how phosphatase inhibition affects overexpressed CK1δ and CK1δ-K38R. Treatment of cells with the phosphatase inhibitor Calyculin A (CalA) triggered rapid phosphorylation and also accumulation of overexpressed CK1δ, indicating that phosphorylated and hence inactive CK1δ is stable (Fig. 2a, left panel). Interestingly, CalA also triggered phosphorylation and accumulation of the kinase-dead CK1δ-K38R, though not to the same level of phosphorylation as CK1δ (Fig. 2a, right panel). These data indicate that CK1δ-K38R is phosphorylated in trans by unknown kinase(s). To further delineate whether CK1δ stabilization depends on kinase inhibition, phosphorylation of the CK1δ C-terminal tail, or phosphatase inhibition, we generated a U2OStx cell line expressing CK1δ-S/A, a mutant kinase wherein all phosphorylation sites in the tail were replaced by alanyl residues. CK1δ-S/A was expressed at a lower level than CK1δ. Upon CalA treatment, CK1δ-S/A remained unphosphorylated, indicating that most of the CalA-induced phosphorylation occurs at the tail. Furthermore, CK1δ-S/A levels decreased suggesting that phosphatase inhibition per se destabilizes CK1δ-S/A. In contrast, upon PF670 treatment, CK1δ-S/A accumulated compared to untreated, indicating that inhibition of enzymatic activity stabilizes the kinase independent of tail phosphorylation (Fig. 2b). Thus, CalA treatment induces tail phosphorylation, which inhibits and thereby stabilizes the kinase.

**Fig. 2.**
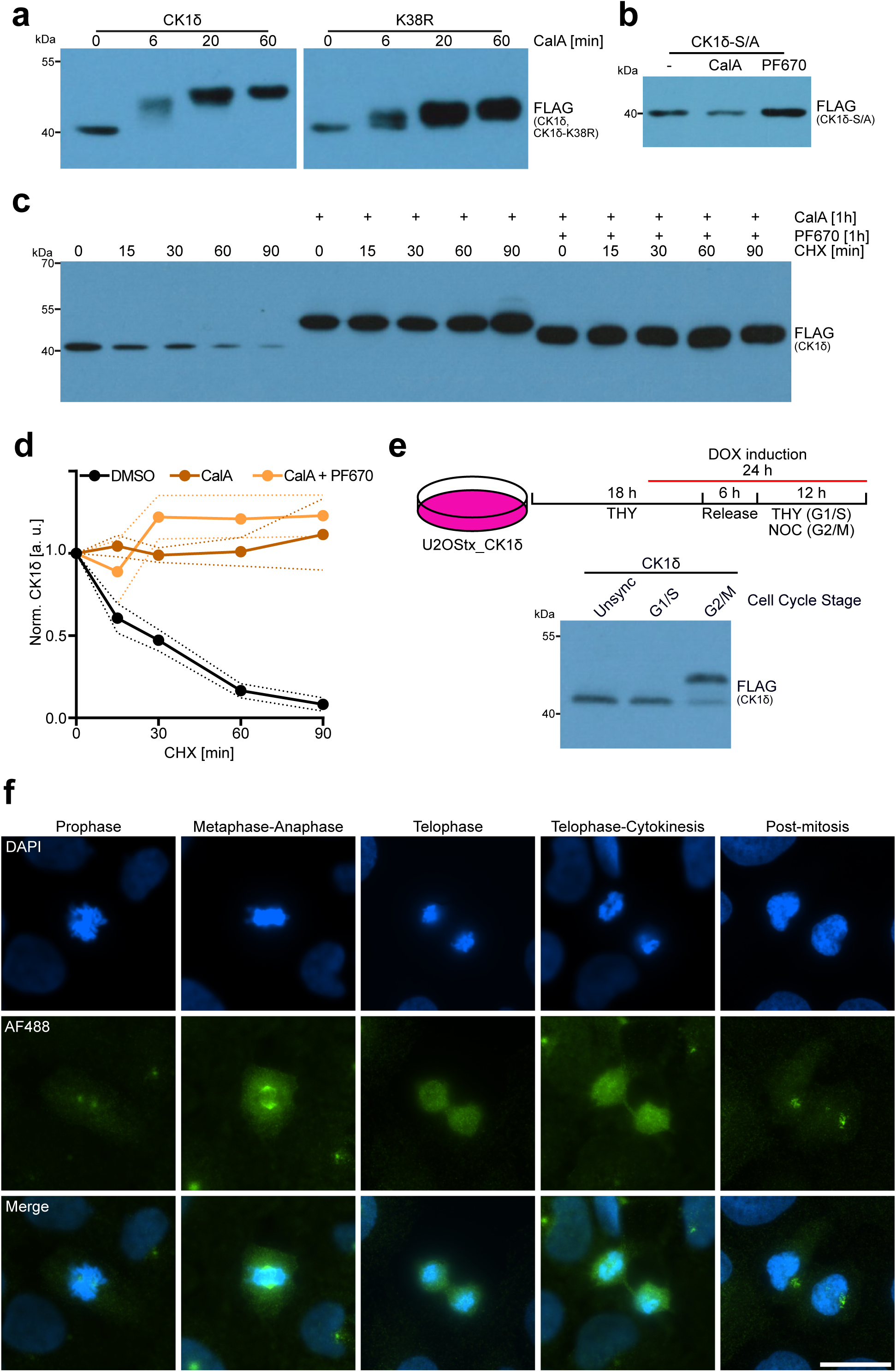
Phosphorylation and inhibition stabilizes CK1δ during G2/M. **a** Phosphatase inhibition by CalA (80 nM) results in (auto)phosphorylation and accumulation of CK1δ and CK1δ-K38R. CK1δ-K38R is phosphorylated to a lesser extent than CK1δ. **b** CK1δ-S/A is destabilized by CalA (80 nM, 1 h) and stabilized by PF670 (1 µM, 1 h). **c** Phosphorylated CK1δ is stable. CK1δ-ovexpressing U2OStx cells were pretreated for 1 h with either CalA alone or with CalA and PF670 followed by a CHX chase and analysis by immunoblotting against the FLAG epitope. **d** Quantification of **c** Results shown are averages of three independent experiments and are represented as the mean ± SD. **e** CK1δ is hyperphosphorylated in cells arrested in G2/M. U2OStx_ CK1δ cells were arrested either in G1/S, G2/M, or unsynchronized (Unsync). Samples were analyzed by immunoblotting against the FLAG epitope. **f** CK1δ is released from centrosomes during late mitosis. Upper panels: DAPI-staining of CK1δ-expressing cells was used to identify stages of the cell cycle. Middle panels. Immunofluorescence analysis of CK1δ with FLAG antibodies. CK1δ is present at the apparent centrosomes in prometaphase (left panel). During metaphase-anaphase, CK1δ is found at the polar microtubules. During telophase and telophase-cytokinesis, CK1δ exhibits a diffuse localization throughout the cell followed by a re-localization to the centrosome in each daughter cell post-mitosis (right panel). Lower panels: Merged DAPI and AF488 image. Scale bar = 20 µm.

We then asked how (auto)phosphorylation affects turnover of overexpressed CK1δ. CK1δ was rapidly degraded when cells were treated with CHX (Fig. 1c, 2c), while CalA treatment resulted in accumulation and stabilization of phosphorylated CK1δ (Fig. 2c, d). Dual treatment with CalA and PF670 triggered partial phosphorylation of CK1δ at a level comparable to that of CK1δ-K38R, suggesting that PF670-inhibited CK1δ was phosphorylated by unknown kinase(s). Because purified recombinant CK1δ fully autophosphorylates *in vitro* without the contribution of other kinases (*12*), these data suggest that in the cell a portion of the CK1δ autophosphorylation sites may be phosphorylated in a redundant manner by unknown kinase(s). The partially phosphorylated PF670-inhibited CK1δ accumulated at elevated levels and was not degraded over the course of a CHX chase (Fig. 2c, d). Hence, partial phosphorylation of CK1δ by unknown kinase(s) as well as complete autophosphorylation were both sufficient to stabilize the enzyme.

Given that CK1δ is involved in the cell cycle and phosphorylated CK1δ does not accumulate in *vivo/cellulo*, we hypothesized that the inhibition and stabilization of CK1δ by (auto)phosphorylation might be transient and of physiological significance during the cell cycle. We therefore arrested cells either in G1/S via a double thymidine block or in G2/M via a thymidine-nocodazole block (*36*). Additionally, we enriched for cells arrested in mid-mitosis by mitotic shake-off (*37*) (Fig. 2e, upper part). In unsynchronized and in G1/S-arrested cells, CK1δ was hypo- or unphosphorylated. However, CK1δ accumulated in its phosphorylated form when cells were arrested at G2/M (Fig. 2e, lower part). The data indicate that (auto)phosphorylated/inhibited CK1δ accumulates during the cell cycle at G2/M, presumably facilitated by modulation of the kinase/phosphatase activity during mitosis.

We then probed for CK1δ via immunofluorescence (IF) in cells at various stages of mitosis to correlate the observed hyperphosphorylated state of CK1δ with its subcellular localization (Fig. 2f). During early mitosis (prophase), CK1δ was enriched in two punctate structures that, consistent with the known localization of CK1δ (*14, 31*), likely represent the duplicated centrosomes. In meta- and anaphase, CK1δ appears to be associated with the polar microtubules as previously described (*20*). Interestingly, in telophase, as chromosomes partitioned to daughter cells, and during cytokinesis CK1δ was diffusely distributed throughout the dividing cell. Finally, post-mitosis, when the nuclear envelope has reformed, CK1δ relocated to the single centrosome in each daughter cell. Phosphorylation-dependent inhibition and thus stabilization of CK1δ therefore appears to be important during mitosis, where it could prevent the degradation of unbound CK1δ such that sufficient kinase is present to reoccupy the centrosomes after cell division.

### Co-expression with PER2 stabilizes CK1δ

Since CK1δ was stable in control cells while overexpressed kinase was rapidly degraded, we examined if co-overexpression of CK1δ binding partners can sequester and thereby stabilize excess kinase. A fraction of CK1δ is found in a complex with PERs and CRYs of the circadian clock (*29*). CK1δ directly interacts with PERs while CRYs bind to and stabilize PERs (*21, 38–40*). Transient co-expression in HEK293T cells of CK1δ with PER2 and/or CRY1 revealed a PER2-dependent stabilization of CK1δ while CRY1 did not stabilize the kinase (Fig. 3a, left part). CK1δ-K38R was more stable in the absence of PER2 compared to wild-type CK1δ, yet CK1δ-K38R was also significantly stabilized by co-expression with PER2 (Fig. 3a, right part). It is worth noting that PER2 was hyperphosphorylated upon co-expression with CK1δ relative to PER2 co-expressed with CK1δ-K38R, as expected, whereas PER2-stabilized CK1δ interestingly remained hypophosphorylated.

**Fig. 3.**
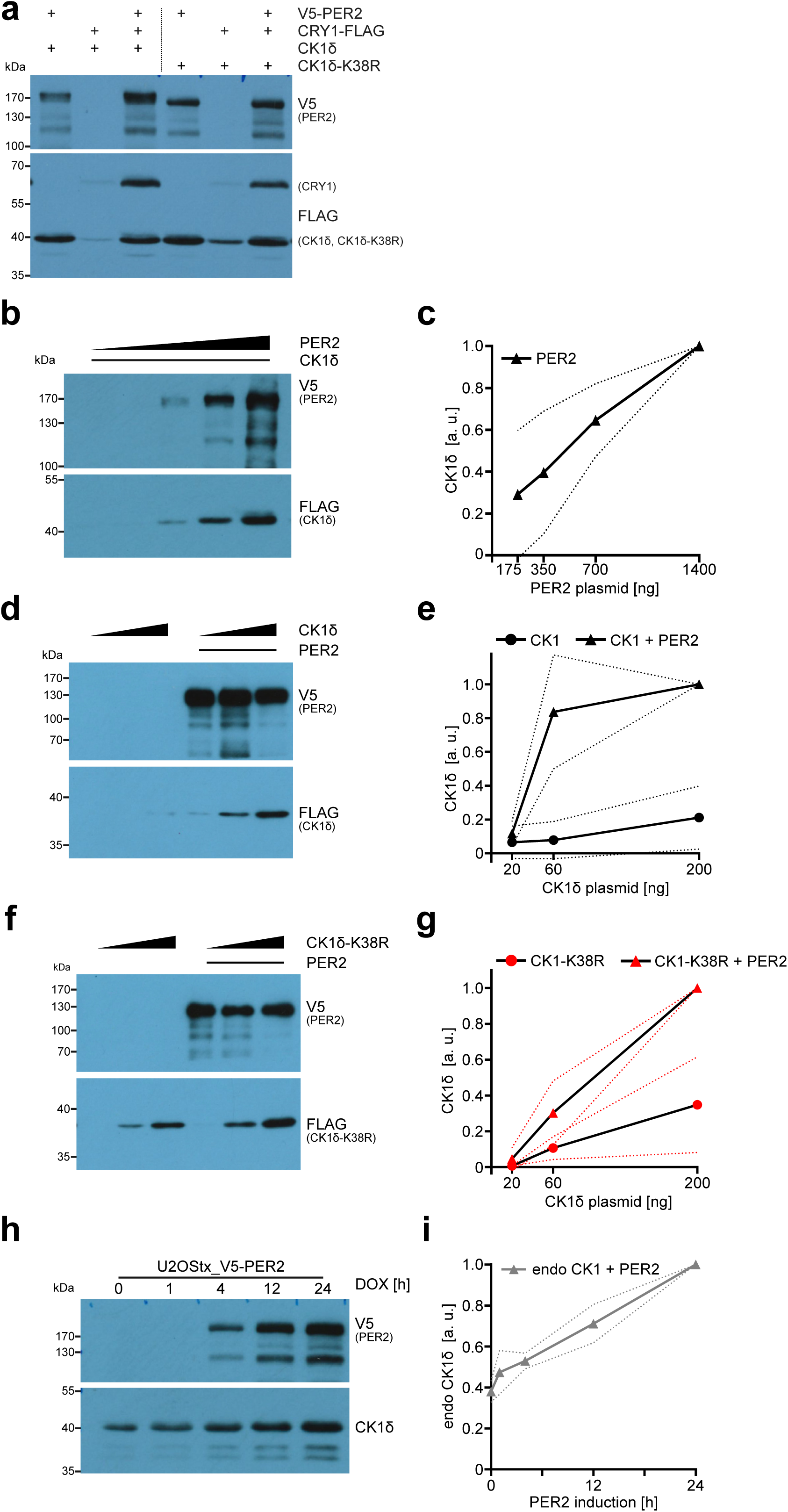
Co-expression of PER2 stabilizes CK1δ. **a** PER2 stabilizes its interaction partners CRY1 and CK1δ whereas CRY1 has no effect on CK1δ. The indicated combinations of V5-tagged PER2, FLAG-tagged CRY1, and either CK1δ or CK1δ-K38R (both FLAG-tagged) were transiently expressed in HEK293T cells for 24 h. The proteins were detected by immunoblotting with the indicated antibodies. Co-expression with PER2 stabilizes CK1δ and to a lesser extent CK1δ-K38R. CRY1 neither stabilizes CK1δ nor CK1δ-K38R. **b** CK1δ is stabilized dependent on PER2 dosage. Co-expression in HEK293T cells of CK1δ (200 ng DNA) with increasing PER2 (175 ng, 350 ng, 700 ng, 1400 ng DNA) results in PER2 dose-dependent stabilization of CK1δ. **c** Quantification of **b**. Results are shown as averages of three independent experiments. Data were normalized to the maximum and represented as the mean ± SD. **d** CK1δ is stabilized dependent on presence of PER2. Co-expression in HEK293T cells of a fixed amount of PER2 (1400 ng DNA) with increasing CK1δ (20 ng, 60 ng, 200 ng DNA) results in CK1δ dose-dependent stabilization of CK1δ. **e** Quantification of **d**. Results are shown as normalized averages of three independent experiments and are represented as the mean ± SD. **f** CK1δ-K38R is stable without PER2 and stabilized by PER2 to a lesser extent than CK1δ. Co-expression in HEK293T cells of a fixed amount of PER2 (1400 ng DNA) with increasing CK1δ-K38R (20 ng, 60 ng, 200 ng DNA). **g** Quantification of **f**. Results are shown as normalized averages of three independent experiments and are represented as the mean ± SD. **h** Endogenous CK1δ co-accumulates with overexpressed PER2. U2OStx_V5-PER2 cells were analyzed at the indicated time points after DOX-induction by immunoblotting against the V5 epitope and endogenous CK1δ. **i** Quantification of **h**. Results are shown as normalized averages of three independent experiments and are represented as the mean ± SD.

When fixed amounts of CK1δ were co-expressed with increasing amounts of PER2, CK1δ was stabilized in a PER2 dose-dependent manner (Fig. 3b, c). Conversely, when PER2 dose was fixed and co-expressed with increasing amounts of CK1δ, we observed PER2-dependent stabilization of the kinase (Fig. 3d, e). CK1δ-K38R was also stabilized when PER2 was co-expressed but to lesser extent due to the inherent stability of CK1δ-K38R even in the absence of PER (Fig. 3f, g). These data show that unassembled active CK1δ is rapidly degraded and that degradation is stimulated by kinase activity.

We then asked whether PER2 would affect the expression of endogenous CK1δ. We generated stable cell lines overexpressing V5 epitope-tagged PER2 (U2OStx_V5-PER2) in a DOX-dependent manner. When PER2 expression was induced, endogenous CK1δ accumulated in parallel with PER2 (Fig. 3 h, i). These data indicate that CK1δ is also expressed by the endogenous locus at somewhat higher levels than needed. Excess orphan CK1δ is degraded but can be stabilized by expression of an interaction partner such as PER2.

### Stabilized CK1δ is predominantly nuclear

Next, we analyzed the subcellular localization of CK1δ by IF. Endogenous CK1δ was concentrated at centrosomes (Fig. 4a, left panels) as previously described (*31, 41*). Otherwise CK1δ was found throughout the cell with a slight enrichment in the nucleus. Overexpressed CK1δ was highly enriched at centrosomes (Fig. 4a, middle panels). Additional overexpressed levels of CK1δ preferentially accumulated in the cytosol. Kinase-inactive CK1δ-K38R was expressed at higher levels than CK1δ, accumulated in the nucleus, in agreement with a previous report (*20*), and did not localize to the centrosomes (Fig. 4a, right panels). The data suggest that the amount and subcellular localization of CK1δ depend on its activity and that excess active CK1δ is preferentially degraded in the nucleus, consistent with the nuclear localization of the CDH1 adaptor of APC/C (*42*).

**Fig. 4.**
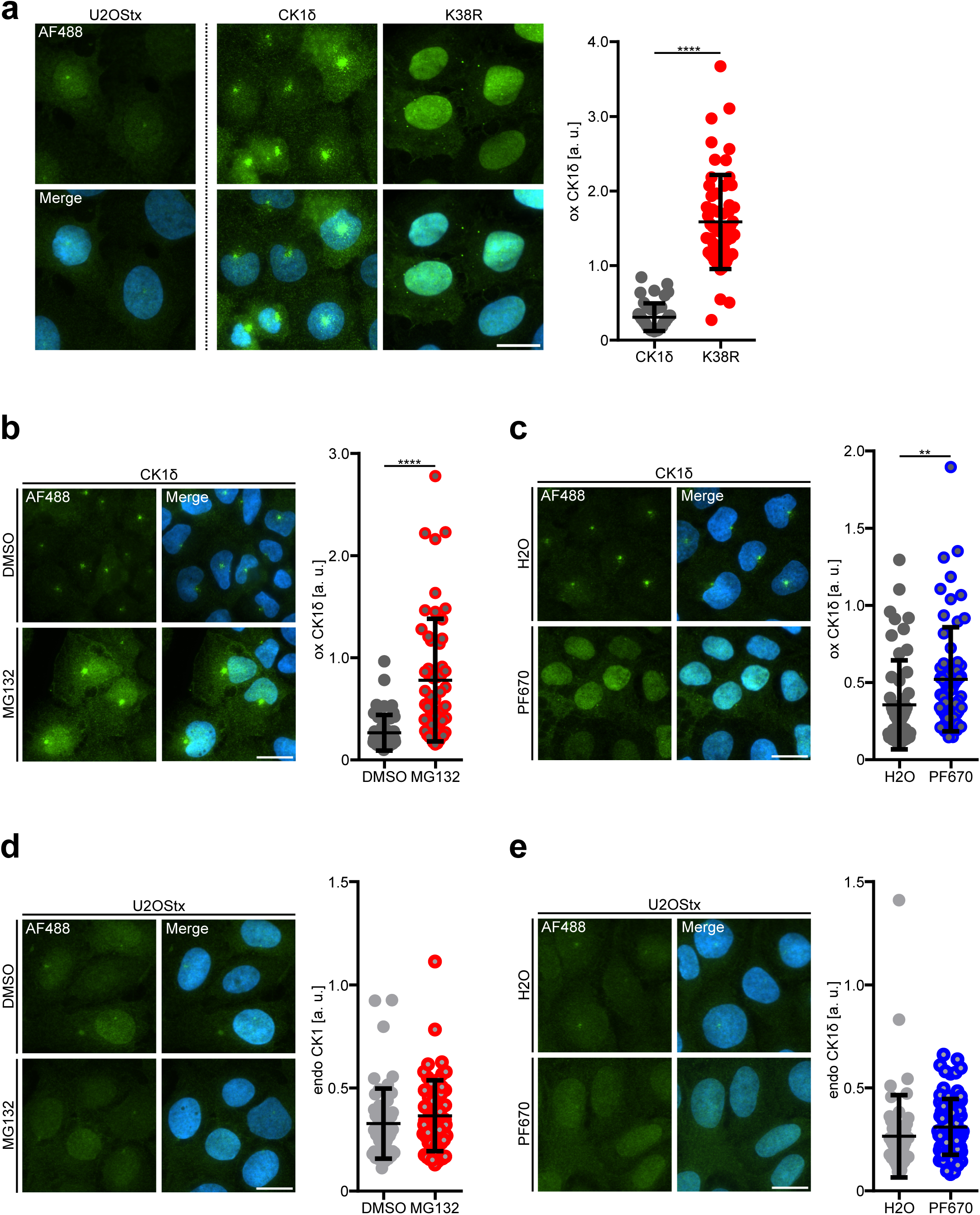
Stabilized CK1δ is predominantly nuclear. **a** Immunofluorescence analysis of the subcellular localization of endogenous CK1δ and overexpressed FLAG-tagged CK1δ and CK1δ-K38R. Left panels: Endogenous CK1δ in U2OStx cells is concentrated at centrosomes and slightly enriched in the nucleus over the cytosol. U2OStx cells were stained with an antibody against CK1δ (upper panel). Middle and right panels: Overexpressed active CK1δ exhibits cytoplasmic and nuclear staining and enrichment at centrosomes. CK1δ-K38R exhibits predominantly nuclear staining with some punctate cytoplasmic staining but no predominant centrosomal localization. U2OStx_CK1δ and U2OStx_CK1δ-K38R cells were induced with DOX for 24 h and stained with an antibody against the FLAG epitope (upper panels). Lower panels show a merge with DAPI staining. Scale bar = 20 µm. Quantification (graph) shows average fluorescence intensities of overexpressed CK1δ (ox CK1δ; black circles) and CK1δ-K38R (red circles) in at least 50 individual cells (n ≥ 50 cells). Data are represented as mean ± SD. *** indicates p-value < 0.001. **b** Proteasomal inhibition by MG132 results in accumulation of CK1δ predominantly in the nucleus. U2OStx_CK1δ cells were DOX-induced for 24 h and treated with 5 µM MG132 or DMSO (vehicle control) for 4 h and then analyzed by IF. Scale bar = 20 µm. Quantification (right panel) shows a significant increase in CK1δ levels in MG132-treated cells. n ≥ 50 cells; mean ± SD; **** indicate p-value < 0.0001. **c** Kinase inhibition by PF670 results in accumulation of CK1δ in the nucleus. U2OStx_CK1δ cells were DOX-induced for 24 h, treated with 1 µM PF670 or H2O (vehicle control) for 4 h, and then analyzed by IF. Scale bar = 20 µm. Quantification shows a significant increase in CK1δ levels in PF670-treated compared to control cells. n ≥ 50 cells; mean ± SD; ** p-value < 0.01. **d** Proteasomal inhibition by MG132 does not affect expression and localization of endogenous CK1δ in U2OStx cells. Scale bar = 20 µm. Quantification shows no significant difference in CK1δ levels in MG132-treated and control cells. n ≥ 50 cells; mean ± SD. **e** PF670 does not affect expression and localization of endogenous CK1δ. Scale bar = 20 µm. Quantification shows no significant difference in CK1δ levels in PF670-treated and control cells. n ≥ 50 cells; mean ± SD.

We then treated CK1δ-overexpressing cells and U2OStx control cells with MG132 or PF670, which both stabilize overexpressed CK1δ (see Fig. 1e, f). Treatment of CK1δ-overexpressing cells with MG132 resulted in approximately 3-fold accumulation of the kinase compared with control cells treated with DMSO, with no apparent difference in subcellular localization (Fig. 4b). Interestingly, centrosomal staining increased upon MG132 treatment of U2OStx cells, indicating that centrosomal binding sites were not saturated. When the CK1δ-overexpressing cells were treated with PF670, CK1δ was stabilized to a lesser extent (1.5-fold) than with MG132. However, the kinase accumulated almost exclusively in the nucleus and centrosomal staining decreased (Fig. 4c) a. These data suggest that inhibition of kinase activity leads not only to stabilization of overexpressed CK1δ but also to its depletion from the cytosol, including centrosomal binding sites, and accumulation in the nucleus. Because inhibition of the proteasome leads to stabilization of overexpressed CK1δ in the cytosol and nucleus, kinase activity appears to be required not only for nuclear degradation of CK1δ but also for its export from the nucleus, consistent with the nuclear accumulation of the catalytically inactive CK1δ-K38R. Treatment of U2OStx control cells with MG132 or PF670 resulted in a small, non-significant increase in CK1δ levels (Fig. 4d and e). Similar to data above, these data suggest that the rate of kinase synthesis in U2OStx control cells is insufficient to significantly increase CK1δ levels within the treatment windows of MG132 and PF670.

### Active orphan CK1δ is primarily degraded in the nucleus

Considering that CK1δ is an APC/C^Cdh1^ client (*33*), that CDH1 is located in the nucleus (*42*), and that CK1δ exhibits nucleocytoplasmic shuttling (*20*), we further analyzed the relationship between subcellular localization and degradation of the kinase. We constructed CK1δ versions that contained either the SV40 nuclear localization signal (NLS) (*43*) or the HIV-1 nuclear export signal (NES) (*44*) at the N-terminus, hereafter referred to as NLS-CK1δ and NES-CK1δ, respectively. U2OStx cells were transiently transfected with NLS-CK1δ and NES-CK1δ expression vectors, respectively. Expression was induced with DOX for 24 h and cells were treated for 4 h with or without MG132 prior to CK1δ detection by IF. NLS-CK1δ was expressed at a low level and exhibited clear nuclear localization (Fig. 5a). MG132 treatment stabilized NLS-CK1δ about 8-fold (Fig. 5a). The data indicate that overexpressed NLS-CK1δ is rapidly degraded by the proteasome in the nucleus. NES-CK1δ was localized in the cytoplasm and MG132 treatment stabilized NES-CK1δ about 1.8-fold (Fig. 5b). Taken together, the data indicate that overexpressed CK1δ is preferentially degraded in the nucleus.

**Fig. 5.**
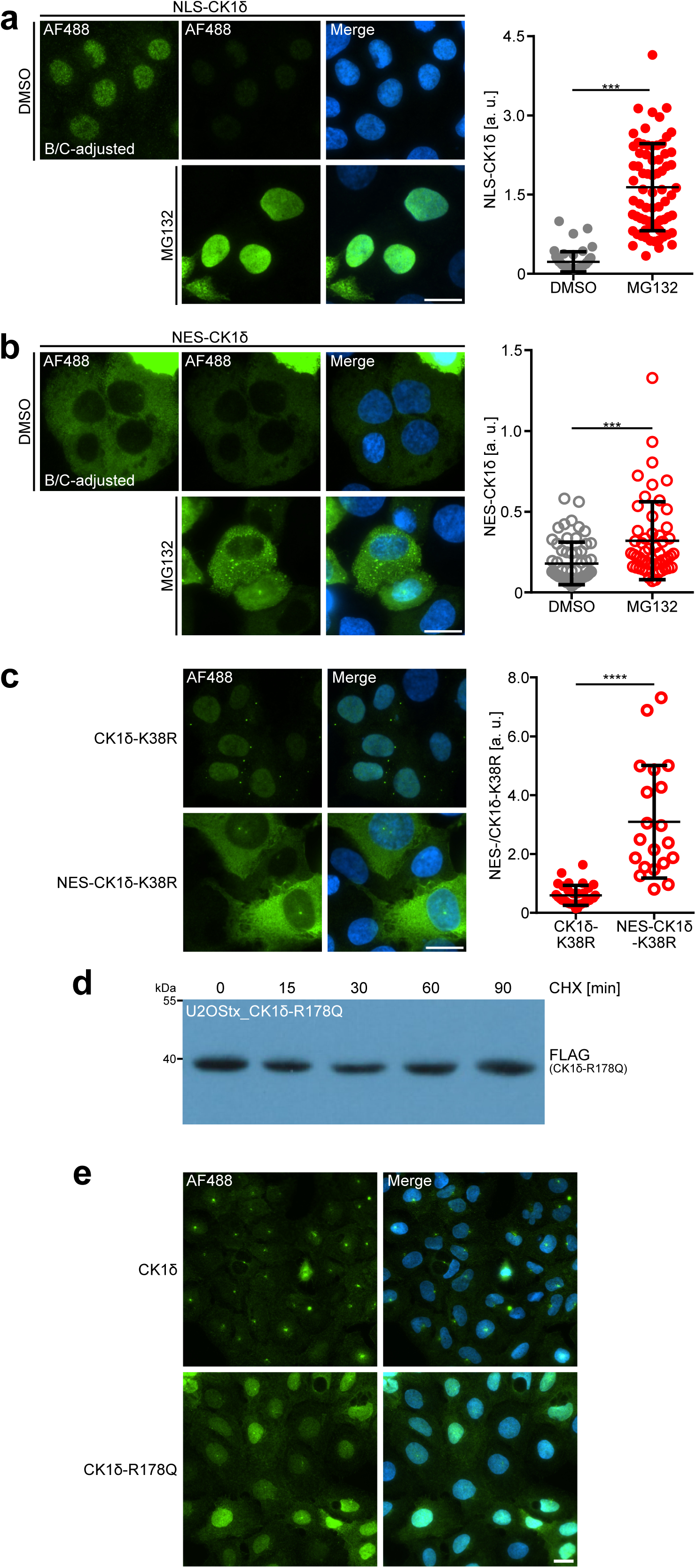
Active CK1δ is primarily degraded in the nucleus. **a** Nuclear localization destabilizes CK1δ. NLS-CK1δ, transiently overexpressed in U2OXts cells, exhibits nuclear localization and accumulates at ∼8-fold higher levels upon proteasomal inhibition by MG132. Left panel: brightness and contrast have been adjusted (B/C-adjusted) to clearly visualize NLS-CK1δ. Middle panels: B/C in upper and lower panels were normalized for comparison of CK1δ levels in DMSO-treated versus MG132-treated cells. Right panels: merge with DAPI stain. Scale bar = 20 µm. Quantification (graph on the right) shows an ∼8-fold increase in NLS-CK1δ upon MG132 treatment. n ≥ 50 cells; mean ± SD. *** indicate p-value < 0.001. **b** Transiently overexpressed NES-CK1δ exhibits cytoplasmic localization and accumulates at a ∼1.8-fold higher level upon proteosomal inhibition by MG132. Left-panel: B/C-adjusted image to clearly visualize NES-CK1δ. Middle panels: B/C in upper and lower panel were normalized for comparison of NES-CK1δ levels in DMSO-treated versus MG132-treated cells. Right panels: merge with DAPI stain. Scale bar = 20 µm. Quantification shows an ∼1.8-fold increase in NES-CK1δ upon MG132 treatment. n ≥ 50 cells; mean ± SD. *** indicate p-value < 0.001. **c** Addition of an NES stabilizes CK1δ-K38R in the cytosol. Top panels: Transiently overexpressed CK1δ-K38R localizes to the nucleus of U2OStx cells. Lower panels: NES-CK1δ-K38R exhibits cytoplasmic localization and increased expression. Scale bar = 20 µm. Quantification indicates ∼6-fold higher expression of NES-CK1δ-K38R in comparison to CK1δ-K38R. n ≥ 30 cells, mean ± SD. **** p-value < 0.0001. **d** Overexpressed CK1δ-R178Q is stable. U2OStx_ CK1δ-R178Q were DOX-induced for 24 h and then subjected to a CHX-chase. Samples were collected at the indicated time points and analyzed by immunoblotting against the FLAG epitope. **e** CK1δ-R178Q is enriched in the nucleus and accumulates at elevated levels. U2OStx_CK1δ cells (upper panels) and U2OStx_CK1δ-R178Q (lower panels) were DOX-induced for 24 h and then analyzed by IF with antibody against the FLAG epitope (left panels). Right panels show a merge with DAPI staining. Scale bar = 20 µm.

To further assess the role of kinase activity on localization and degradation, we generated a kinase-dead CK1δ-K38R version harboring the HIV-1 NES (NES-CK1δ-K38R). CK1δ-K38R and NES-CK1δ-K38R were transiently expressed for 24 h in U2OStx cells and then detected by IF. CK1δ-K38R localized primarily to the nucleus (Fig. 5c, upper panels), similar to the localization of CK1δ-K38R expressed from a stably transformed cell line (see Fig. 4a). In contrast, NES-CK1δ-K38R localized to the cytoplasm (Fig. 5c, lower panels), indicating that the NES was functional. NES-CK1δ-K38R was expressed at a 5-fold higher level than CK1δ-K38R (Fig. 5c). The data suggest that even enzymatically inactive CK1δ is relatively unstable in the nucleus.

Overall, the data indicate that overexpressed active orphan CK1δ is unstable in the nucleus but stable in the cytosol. Enzymatic activity of CK1δ is required for its nuclear export and furthermore accelerates the nuclear degradation of unassembled CK1δ.

### CK1δ with a Tau-like mutation accumulates at elevated levels in the nucleus

The so-called tau mutation is a R178C substitution in CK1ε that reduces kinase activity and is associated with a short circadian period (*45–47*). Since our data showed that kinase activity is required for nuclear degradation and export of CK1δ, we examined expression and localization of CK1δ with a Tau-like R178Q substitution. We have recently shown that CK1δ-R178Q is also less active, and U2OStx cells expressing the mutant kinase exhibit a tau-like short period (*12*). When protein synthesis in U2OStx_CK1δ-R178Q cells was inhibited with CHX, the overexpressed orphan CK1δ-R178Q was stable (Fig. 5d). Immunofluorescence analysis showed that CK1δ-R178Q accumulated preferentially in the nucleus and was expressed at a higher level than CK1δ in control cells (Fig. 5e). Therefore, altered expression levels and subcellular localization of CK1δ-R178Q (and possibly CK1ε^Tau^) might contribute to their short-period phenotype in addition to the reduced kinase activity (see Discussion).

## Discussion

CK1δ is a ubiquitously expressed monomeric kinase implicated in a wide variety of cellular pathways and functions. It is inhibited by (auto)phosphorylation of its unstructured C-terminal tail, but in the cell CK1δ is dephosphorylated (*6*), raising the question of how kinase activity is controlled.

Here we characterized posttranslational regulation that contributes to the homeostasis of CK1δ expression. Our data indicate that virtually all detectable CK1δ is assembled and the pool of free kinase is low. In search for interaction partners, unassembled CK1δ shuttles between cytosol and nucleus and is susceptible to rapid degradation by the proteasome (Figure 6).

**Fig. 6.**
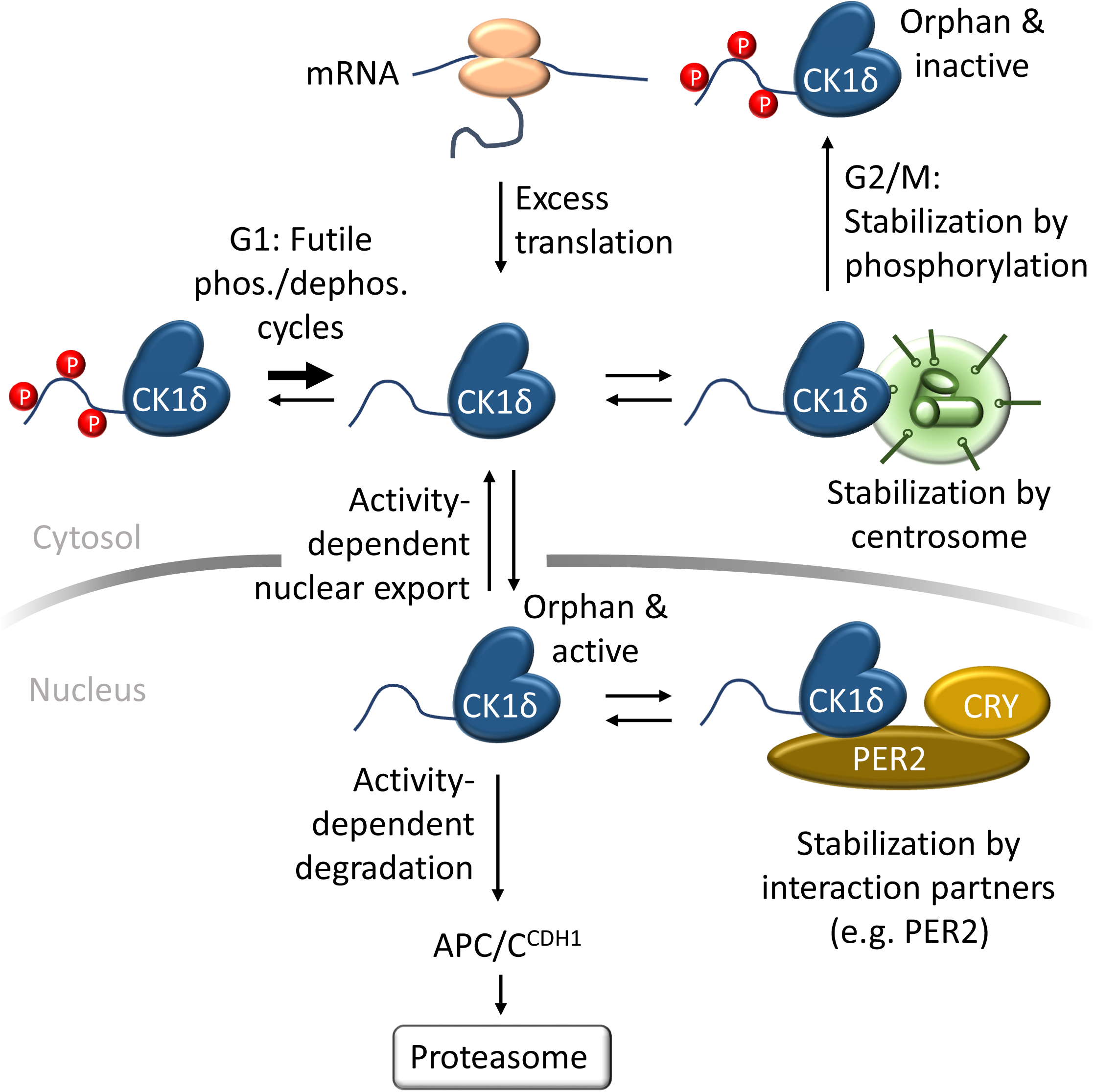
Model of CK1δ homeostasis CK1δ is translated at higher levels than needed and undergoes in G1 futile cycles of autophosphorylation/inhibition and dephosphorylation/activation in the cytosol and/or nucleus. Orphan active CK1δ shuttles between cytosol and nucleus in search of interaction partners and is highly susceptible to proteasomal degradation in the nucleus mediated by APC/C^Cdh1^. Kinase activity facilitates two competing processes: accelerated degradation of unassembled CK1δ in the nucleus and its export from the nucleus, resulting in its stabilization in the cytosol. Interaction of CK1δ with clients such as centrosomes or PER2 stabilizes the kinase. During the cell cycle, CK1δ is released from centrosomes in G2/M. The orphan kinase is not degraded but inactivated and stabilized by (auto-) phosphorylation leaving enough kinase to reassociate with interaction partners after mitosis.

Degradation of orphan, unassembled subunits of protein complexes is a rather common mechanism of quality control ensuring protein homeostasis (*48, 49*). Proteins orphaned by overexpression or mislocalization, are usually committed to proteasomal degradation by dedicated E3 ubiquitin ligases such as Tom1/HUWE1, UBE2O, Not4, Teb4, UBR1, which often recognize degrons that are buried and not accessible when their clients are correctly assembled (*48, 49*).

Our results together with published data show that orphan CK1δ is not degraded by this pathway. It has been previously shown that CK1δ is a client of the APC/C ubiquitin ligase with the adaptor FZR1/CHD1, which recognises two D-boxes in the kinase domain of CK1δ (*33*). APC/C^Cdh1^ is the major degradation pathway because mutation of D-boxes or downregulation of CDH1 prevents CK1δ degradation (*33*). We show here that CK1δ is degraded predominantly in the nucleus, consistent with the reported nuclear localization of FZR1/CDH1 (*42*) and the activity of APC/C^Cdh1^ in G1 phase of the cell cycle (*33, 50*). We also show that APC/C^Cdh1^ specifically targets orphan CK1δ. Although CDH1 recognizes CK1δ through its D-boxes (*33*), we additionally show that rapid degradation of orphan CK1δ is dependent on kinase activity. Therefore, our data suggest that CK1δ either phosphorylates APC/C^Cdh1^, which may improve CK1δ recognition, or an unknown factor, which then commits the kinase to degradation via APC/C^Cdh1^. We also show that export of CK1δ from the nucleus depends on kinase activity but is independent of autophosphorylation, suggesting that CK1δ phosphorylates and activates a factor required for export. Thus, kinase activity controls two competing processes: binding to CDH1, which leads to degradation in the nucleus, and export from the nucleus to the cytosol, which stabilizes the kinase. Hence, mutations compromising kinase activity of CK1δ could therefore affect phosphorylation of clients not only directly but also indirectly via stabilization and accumulation of unassembled kinase in the nucleus (Figure 6).

We show that the mutant CK1δ-R178Q was stable and accumulated in the nucleus. CK1δ-R178Q is a model of CK1ε^Tau^ (R178C). Both mutant kinases are associated with a short circadian period and have severely reduced kinase activity toward primed sites, prephosphorylated at the -3 position and, to a lesser extent, toward unprimed sites and (*12, 45, 46*). Period shortening by CK1ε^Tau^ was attributed to slower priming-dependent phosphorylation of the inhibitory FASPS region in PER2, which in turn promotes accelerated priming-independent phosphorylation of the ß-TrCP phosphodegron despite the reduced kinase activity of CK1ε^Tau^ (*11, 26, 51*). However, Tau hamsters also express fully active CK1δ and it is not known why the less active CK1ε^Tau^ is dominant over CK1δ, which is considered the major kinase of PERs (*5, 29*). Our data strongly suggest that unassembled CK1ε^Tau^, being stable and accumulating in the nucleus, can outcompete free, unstable CK1δ in search for PERs.

CK1δ expressed in U2OStx cells by the endogenous gene accumulated after 24 h when its binding partner PER2 was overexpressed but did not detectably accumulate upon treatment of cells for 4 h with PF670. This suggests that although stabilization of kinases by binding partners has a competitive advantage over degradation, the rate of synthesis of endogenous CK1δ is so low that it takes a long time for stabilized kinase to accumulate detectably. Thus, transcription of *CSNK1D* appears to be adapted to cellular requirements and may therefore be regulated.

Among the various CK1δ binding sites, the centrosome is a uniquely identifiable structure and therefore readily provides evidence for CK1δ binding. Centrosomal binding of CK1δ increased when the kinase was overexpressed/stabilized in U2OStx cells, indicating that binding was not saturated. We show that the export-deficient CK1δ-K38R accumulated in the nucleus but not on centrosomes, whereas the cytosolic NES-CK1δ-K38R version accumulated on centrosomes. Thus, centrosomal accumulation of CK1δ is dynamic and relies on multiple rounds of nucleo/cytoplasmic shuttling (Figure 6).

CK1δ is kept dephosphorylated by futile cycles of (auto)phosphorylation and dephosphorylation (*6*). We found that PER2-bound CK1δ remained dephosphorylated, whereas PER2 was hyperphosphorylated. This suggests that PER2-bound CK1δ either cannot autophosphorylate or that CK1δ is a much better phosphatase substrate than PER2.

Finally, we found that CK1δ was efficiently (auto)phosphorylated and thus inactive in U2OStx cells arrested in G2/M during the cell cycle. At late stages of mitosis, CK1δ appeared to be orphaned but stable as it accumulated in larger amounts and was distributed throughout the cell. Therefore, the physiological function of autophosphorylation might be to stabilize unassembled CK1δ to escape degradation during late mitosis when its ubiquitin ligase, APC/C^Cdh1^, becomes active again (*50*). Apparently, the phosphatase(s) that keep CK1δ dephosphorylated in G1 are not active during mitosis (Figure 6). Given the low rate of endogenous CK1δ synthesis, protecting orphan CK1δ from degradation during mitosis would maintain a sufficiently large kinase pool that could be rapidly reactivated in early G1 phase and bind to centrosomes and other partners, including PER proteins of the circadian clock. Because the cell cycle and the circadian clock are functionally linked and influence each other (*52–55*), the inactivation of CK1δ during mitosis could also be part of a mechanism by which the cell cycle acts on the circadian clock.

In summary, our data show that homeostasis of the apparently simple monomeric CK1δ is regulated in a complex manner by a posttranslational mechanism involving dynamic stabilization by interacting partners and activity-dependent subcellular shuttling and nuclear degradation. Our data also suggest that autophosphorylation of CK1δ provides a specific mechanism for inactivation and thus stabilization of orphan kinase during mitosis.

## Materials and Methods

### Cell culture

T-REx-U2OS (U2OStx; Life Technologies) and HEK293T (ATCC) cells were maintained in Dulbecco’s Modified Eagle Medium (DMEM) supplemented with 10% Fetal Bovine Serum (FBS) and 1x Penicillin-Streptomycin. Cell culture reagents were obtained from Life Technologies. Cells were grown and maintained at 37 °C in a humidified incubator containing 5% CO_2_.

### Plasmid constructs

pcDNA4/TO-derived vectors were constructed by cloning the coding regions of human casein kinase 1 isoform delta (CK1δ), murine Period circadian protein homolog 2 (mPer2), and murine cryptochrome 1 (mCry1) downstream of the CMV-TetO2 promoter which allows for doxycycline-inducible protein expression in cells harboring the tetracycline-repressor cassette (tx; i.e. U2OStx) and constitutively high expression in cells lacking the tetracycline-repressor cassette (i.e. HEK293T). Gene synthesis (GenScript) and ligation-free overlap cloning(*56*) was used to introduce the CK1δ-K38R and CK1δ-R178Q point mutations, assemble the CK1δ-S/A mutant construct, as well as the NLS-CK1δ, NES-CK1δ, and NES-CK1δ-K38R constructs. All plasmids were verified by sequencing (LGC Genomics).

### Generation of stable U2OStx cell lines

U2OStx cells were seeded in a 24-well plate and grown to confluence overnight. Cells were transfected with pcDNA4/TO vectors containing the desired alleles of FLAG epitope-tagged CK1δ, CK1δ-K38R, CK1δ-R178Q, CK1δ-S/A and V5 epitope-tagged mPer2. Stable transfectants were selected by growing cells to sub-confluence in complete media supplemented with 50 µg/mL hygromycin (Invitrogen) and 100 µg/mL zeocin (Invitrogen) over the course of two weeks. Resulting single cell colonies after selection were isolated for further experimentation as previously described (*54, 57*). Unless otherwise stated, induction of protein expression was typically performed by addition of 10 ng/mL doxycycline (DOX; Thermo Fisher Scientific CAT# 631311) 24 h prior to downstream experiments.

### Quantitative PCR

Cells were grown in 24-well plates (p24) and induced with DOX for 24 h. Cells were then washed with PBS, and RNA was extracted by adding 200 µL of peqGOLD TriFast Reagent per p24 well. Succeeding RNA extraction steps were done following manufacturer’s instructions. cDNA was reverse-transcribed from 500 ng total RNA extract with Maxima Reverse Transcriptase (Thermo Fisher Scientific) following manufacturer’s protocols. Quantitative PCR (qPCR) was performed on a StepOne™ Real-Time PCR System (Applied Biosystems, Themo Fisher Scientific, CAT# 4376598) using Maxima SYBR Green qPCR Master Mix (Thermo Fisher Scientific) following manufacturer’s protocols. The following primer pairs were used to target CK1δ and GAPDH, respectively:

CK1-F: AACCAAACACCCTCAGCTCCAC CK1-R: GCCCCAGCAGCTCCATCACCAT GAPDH-F: TGCACCACCAACTGCTTAGC GAPDH-R: ACAGTCTTCTGGGTGGCAGTG

Relative CK1δ gene expression was normalized to GAPDH and calculated using the ΔΔC_T_ method. Further normalization to the DOX-induced U2OStx sample was done to determine fold induction of gene expression.

### Protein extraction, SDS-PAGE, and Immunoblotting

Protein extraction was performed by scraping the cells from the culture dish and spun down to collect the cell material. After removing the spent media on ice, cells were treated with ice-cold lysis buffer (25 mM Tris-HCl, pH 8.0, 150 mM NaCl, 0.5% Triton X100, 2 mM EDTA, 1 mM NaF with freshly added protease inhibitors 3 µg/mL PMSF, 10 µg/mL Leupeptin, 34 µg/mL PepstatinA) for 5 min on ice. Resulting crude lysates were then sonicated for 5 min in an ultrasonic bath (Merck) for 5 min. Lysates were pre-cleared from cell debris by centrifugation at 16,000 g for 15 min at 4°C. Protein concentration was quantified photometrically (NP80, Implen). Unless otherwise stated, protein extracts were typically loaded in equal amounts of 50 µg onto PAGE gels. SDS-PAGE and Immunoblotting was performed as described(*12*).

### Cycloheximide chase; MG132, PF670, and CalA treatment

To arrest protein translation for a cycloheximide (CHX) chase, CHX was added to cells to a final concentration of 10 µg/mL. Untreated samples were taken as the 0 min CHX sample. Samples were then taken post-CHX for protein extraction and immunoblotting as indicated.

To inhibit the proteasome, cells were treated with 20 µM MG132 (Abmole Bioscience CAT# AMO-M1902) for 4 h prior to immunoblotting; 4 µM MG132 for 4 h prior to immunofluorescence. To inhibit CK1δ; cells were treated with 1 µM PF670462 (Tocris Biosciences CAT# 3316) for 4 h prior to immunoblotting and immunofluorescence. To inhibit phosphatases, cells were treated with 80 nM CalA (LC Labs Cat# C-3987) for 0 min (untreated), 6, 20, and 60 min. All chemical treatments were performed with the corresponding solvent (vehicle) control: DMSO for MG132 and CalA, and H_2_O for PF670.

### Cell cycle arrest

Cell cycle arrest protocol was adapted from Apraiz et al., 2017 (*36*). U2OStx_CK1 was seeded onto four 10 cm dishes to 50% confluence. For the unsynchronized sample, no additional treatment with thymidine or nocodazole was performed. For both the G1- and G2-arrested samples, after 24 h post-seeding, cells were treated with thymidine (THY; final concentration of 2 mM) for 18 h followed by a release into normal media for 6 h. To arrest cells in G1, one 10 cm dish was used and the cells were treated with a second thymidine block for 12 h. To arrest cells in G2, two 10 cm dishes were used and the cells were treated with nocodazole (NOC, final concentration of 50 ng/mL) for 12 h. For the G2/M-arrested cells, an additional mitotic shake-off was performed to enrich the sample for cells in mid-mitosis that are loosely attached to the culture surface(*37*). To induce CK1δ, DOX induction was performed on all samples 24 h prior to the end of the thymidine-nocodazole treatments.

### Protein overexpression in HEK293T cells

HEK293T cells were seeded onto 6-well plates (p6) and transfected with polyethylenimine (PEI) according to the conventional protocol. Briefly, depending on the plate format, up to 1600 ng of plasmid DNA was complexed with 60 µL of PEI (19 mM) for a 10 cm dish; and 200 ng of plasmid DNA was complexed with 8 µL of PEI (19 mM) for a p6. Plasmid-PEI complexation was allowed by incubating at room temperature for at least 20 min. For the PER-CRY-CK1 co-expression experiment, PER and CRY dose was set at 700 ng each whereas the CK1 and CK1-K38R dose was set at 200 ng each. For PER titration, CK1 was set at 200 ng; PER dose was titrated at 0, 175, 350, 700, and 1400 ng. For CK1 titration, PER was set at 1400 ng; CK1 dose was titrated at 20, 60, and 200 ng. For experiments wherein plasmid titration was performed, the remaining plasmid dosage was compensated with empty vector (pcDNA4/TO). Cells were allowed to express protein until 24 h post-transfection wherein the cells were lysed for immunoblotting.

### Immunofluorescence

Cells were seeded on a 24-well plate on 12 mm #1.0 glass coverslips (VWR, CAT# 631-1577P) in 500 µL media. Whenever applicable, transient transfection and/or DOX induction was performed. Immunofluorescence was done 24 h post-induction. Cells were washed one time with PBS and fixed with 4% paraformaldehyde for 10 min at room temperature. Cells were permeabilized with PBS + 0.1% Triton X-100 for 10 min, then washed three times and blocked with PBS + 5% heat-inactivated FBS for 1 h. Cells were then treated with antibodies against FLAG (Monoclonal ANTI-FLAG® M2 antibody produced in mouse; Merck CAT# F3165; 1:200 dilution) or CSNK1D (Anti-Casein Kinase 1 delta/CSNK1D antibody [AF12G4]; abcam CAT# ab85320; 1:1000 dilution) in PBS + 3% BSA + 0.25% Tween20 for 1 h at 37 °C, washed three times with PBS, and then treated with AlexaFluor488-conjugated secondary antibody (αmouse IgG Alexa Fluor™ 488; Thermo Fisher Scientific CAT #A-11001; 1:500 dilution) in PBS + 3% BSA + 0.25% Tween20 for 1 h at 37 °C. Cells were again washed three times, then mounted onto glass slides with ProLong™ Glass Antifade Mountant with NucBlue (Thermo Fisher Scientific) and sealed with nail polish.

### Microscopy

For microscopy, a Nikon Ni-E microscope was used (Nikon Imaging Center, Heidelberg) at a 60X oil immersion objective (Objective: Nikon Plan Apo λ 60x NA 1.40 Oil, Camera: DS-Qi2 black and white camera). The (DAPI, FITC FILTER DETAILS) DAPI and FITC filter sets (DAPI: excitation 390/18, dichroic mirror 416, emission 460/60; FITC: excitation 472/39, dichroic mirror 495, emission 520/35) were used. Exposure times were optimized within each experiment to obtain comparable fluorescence intensities. For image acquisition, the central focal plane was determined and images were taken at 0.3 µm z-steps (11 steps with the 6^th^ step as the central focal plane).

### Image Analysis

Image analysis was done in FIJI (*58*). For immunoblotting images, lanes were defined and the densitometric analysis was performed per lane. For the CHX chase experiments, resulting quantification values were normalized against the starting quantification at the start of the CHX chase. For the HEK293T co-expression experiments, resulting quantification values were normalized against the highest quantification at the highest PER2/CK1δ dose or DOX induction. For each plot, three independent experiments were performed and the data are presented as mean ± SD, asterisks indicate p-value.

For immunofluorescence images, to quantify the AF488 signal, the FITC channel was isolated and used to generate a Z-projection image taking the maximum intensity from each z-plane. Using this Z-projected image, we performed threshold adjustment to create masks which we then used to analyze single cells as particles. Measurements were set to quantify area, mean gray value, and integrated intensity as well as shape descriptors. For each plot, at least 30 cells were measured and the data are presented as mean ± SD, asterisks indicate p-value.

### Quantification and Statistical Analysis

Data were plotted on GraphPad 6. To test for differences between two samples, student’s t-test was performed. To test for difference between three or more samples, ANOVA was performed.

## Author contributions

Conceptualization: MB, ACRD; Methodology: FES, BR, DM, ACRD; Investigation: FES, BR, DM; Supervision: MB, ACRD; Writing. MB, FES

## Data and materials availability

Further information and requests for reagents should be directed to by the lead contact, Michael Brunner. Plasmids and strains generated in this study will be distributed upon reasonable request and corresponding MTA. All other data are available upon request. All data are available in the main text or the supplementary materials.

